# Predicting SARS-CoV-2 epitope-specific TCR recognition using pre-trained protein embeddings

**DOI:** 10.1101/2021.11.17.468929

**Authors:** Youngmahn Han, Aeri Lee

## Abstract

The COVID-19 pandemic is ongoing because of the high transmission rate and the emergence of SARS-CoV-2 variants. The **P272L** mutation in SARS-Cov-2 **S**-protein is known to be highly relevant to the viral escape associated with the second pandemic wave in Europe. Epitope-specific T-cell receptor (TCR) recognition is a key factor in determining the T-cell immunogenicity of a SARS-CoV-2 epitope. Although several data-driven methods for predicting epitope-specific TCR recognition have been proposed, they remain challenging owing to the enormous diversity of TCRs and the lack of available training data. Self-supervised transfer learning has recently been demonstrated to be powerful for extracting useful information from unlabeled protein sequences and increasing the predictive performance of the fine-tuned models in downstream tasks.

Here, we present a predictive model based on Bidirectional Encoder Representations from Transformers (BERT), employing self-supervised transfer learning, to predict SARS-CoV-2 T-cell epitope-specific TCR recognition. The fine-tuned model showed notably high predictive performance for independent evaluation using the SARS-CoV-2 epitope-specific TCR CDR3β sequence datasets. In particular, we found the proline at position 4 corresponding to the **P272L** mutation in the SARS-CoV-2 **S**-protein_269-277_ epitope (**YLQPRTFLL**) may contribute substantially to TCR recognition of the epitope through interpreting the output attention weights of our model.

We anticipate that our findings will provide new directions for constructing a reliable data-driven model to predict the immunogenic T-cell epitopes using limited training data and help accelerate the development of an effective vaccine in response to SARS-CoV-2 variants.

## Introduction

The global population is currently suffering from a pandemic of the coronavirus disease 2019 (COVID-19) caused by the novel coronavirus known as severe acute respiratory syndrome coronavirus 2 (SARS-CoV-2). Since the World Health Organization (WHO) declared COVID-19 as a pandemic on March 11, 2020, there have been 237,251,035 confirmed cases worldwide, and 4,842,805 deaths (GISAID, https://www.gisaid.org/epiflu-applications/global-cases-covid-19/) [1]. Despite the number of vaccinations exceeding 6.4 billion, the pandemic is ongoing because of the high transmission rate and the emergent SARS-CoV-2 variants associated with disease severity and viral escape of humoral immunity [2]. To end the pandemic, many countries and global scientific communities are developing effective vaccines and appropriate treatments in response to these variants.

In addition to the virus-neutralizing antibodies produced by B-cells, cytotoxic CD8^+^ T-cells and the helper CD4^+^ T-cells are essential for viral clearance. T-cells circulating in the blood lead the first response to any virus in the adaptive immune system: they detect infected cells and mount an immune response or directly clear the infected cells, often before symptoms appear [3-6]. The development of effective COVID-19 vaccines, therefore, depends on the identification of T-cell epitopes that can induce T-cell immune responses.

Peptide-major histocompatibility complexes (MHCs) on the cell surface are recognized by T-cells via a dimeric surface protein, the T-cell receptor (TCR), consequently leading to T-cell activation and proliferation by clonal expansion [7]. TCR recognition of a T-cell epitope is therefore crucial for determining the immunogenicity of the epitope. TCRs are generated by genomic rearrangement of the germline TCR loci from a large collection of variable (V), diversity (D), and joining (J) gene segments. During T cell development, most TCRs are formed by a pair of α- and β-chains (90-95% of T cells) via the V(D)J recombination in each locus independently. This rearrangement is estimated to generate 10^18^ different TCRs, providing an enormous diversity of epitope-specific T-cell repertoires [8, 9].

Despite this TCR diversity, recent studies have found that TCRs recognizing a specific target epitope often share common sequence features. Glanville *et al*. [10] and Dash *et al*. [11] have shown a clear signature of the amino acid motif in the complementarity-determining region 3(CDR3) of TCRβ and TCRα that interacts with specific peptides presented by specific MHC molecules. Furthermore, concerted data collection efforts [12-15] and advances in high-throughput TCR sequencing technologies have demonstrated T-cell specificity [16, 17], allowing the development of a data-driven model for predicting epitope-specific TCR recognition [18]. Several methods using position-specific scoring matrices [10], Gaussian processes [19], random forests [20], convolutional neural networks [21], deep generative models [22, 23], and natural language process (NLP)-based deep learning models [24] have been proposed. However, increasing the predictive power of a machine-learning (or deep learning) model remains challenging because of the scarcity of training data: as of October 2019, the VDJdb [15] and McPAS-TCR [13] databases contained about 20,000 and 55,000 epitope-specific TCR sequences, respectively.

Recent advances in NLP have demonstrated that self-supervised learning can be a powerful tool for extracting useful information from unlabeled sequence data [25-27]. One successful approach, Bidirectional Encoder Representations from Transformers (BERT) [26], is a language model pre-trained using a huge amount of unlabeled text data via two self-supervised tasks, masked token prediction and next sentence prediction. BERT models, fine-tuned using a small number of datasets, have shown ground-breaking results in 11 NLP downstream tasks. The self-supervised transfer learning strategy constructs the final model by fine-tuning the self-supervised pre-trained model on a large amount of unlabeled data, using a small amount of labeled data in a downstream task; this strategy can be useful for increasing the predictive power of a deep learning model when there is scarce training data. The self-supervised transfer learning has been demonstrated to help learn protein sequence patterns [28-30]. The Tasks Assessing Protein Embeddings (TAPE) [28] model was pre-trained on 31 million unlabeled protein sequences derived from the Pfam database [31] via two protein-specific self-supervised tasks, amino acid contact prediction and remote homology detection. The TAPE pre-trained model is helpful for improving the predictive performance in supervised downstream tasks such as secondary structure prediction, amino acid contact prediction, remote homology detection, fluorescence landscape prediction, and protein stability landscape prediction. BERTMHC [32], a deep learning model generated by fine-tuning the pre-trained TAPE model, has shown a reliable performance in predicting both peptide-MHC-II binding and presentation, using relatively little training data.

Many sequence-based methods for modeling epitope-specific TCR recognition have used a multiple sequence alignment (MSA) of TCR sequences to identify position-specific amino acid motifs. This makes it difficult to find the critical amino acid positions in both the epitope and the TCR sequence, which can be highly relevant in TCR recognition [10, 11, 22-24, 37]. A recent study [33] of protein language models has shown that the output attentions of BERT-based protein models can capture biologically relevant protein properties. An attention-based deep learning model for peptide-MHC-I binding predictions [34] has shown that the attentions learned by the predictive model can capture critical amino acid positions of the peptides, which help stabilize the peptide-MHC-I bindings.

Here, we present a BERT-based model employing self-supervised transfer learning for predicting SARS-CoV-2 T-cell epitope-specific TCR recognition. The predictive model was generated by fine-tuning the pre-trained TAPE model using epitope-specific TCR CDR3β sequence datasets. The fine-tuned model showed markedly high predictive performance in the independent evaluation using SARS-CoV-2 epitope-specific CDR3β sequence datasets and outperformed the recent Gaussian process-based method. In particular, we found the critical amino acid positions of both epitope and CDR3β sequences, which potentially contribute greatly to the TCR recognition of an epitope, can be captured using the output attention weights of our model. We anticipate that our findings will provide new directions for constructing a reliable model for predicting the immunogenic T-cell epitopes using limited training data and help accelerate the development of an effective vaccine in response to SARS-CoV-2 variants, by identifying potential amino acid motifs highly relevant to the epitope-specific TCR recognition.

## Materials and Methods

### Training process and model architecture

Figure 1 is a schematic representation of the training process of the proposed model. The initial model is cloned from the pre-trained BERT-based TAPE model, adding a classification layer at the end. First, the initial TAPE model is fine-tuned using general epitope-specific CDR3β sequence data, while freezing the embedding layer and top two encoding layers. Next, the final model is fine-tuned from using SARS-CoV-2 epitope-specific CDR3β sequence data derived from Immune Epitope Database (IEDB), while freezing the embedding layer and top six encoding layers.

**Figure 1.**
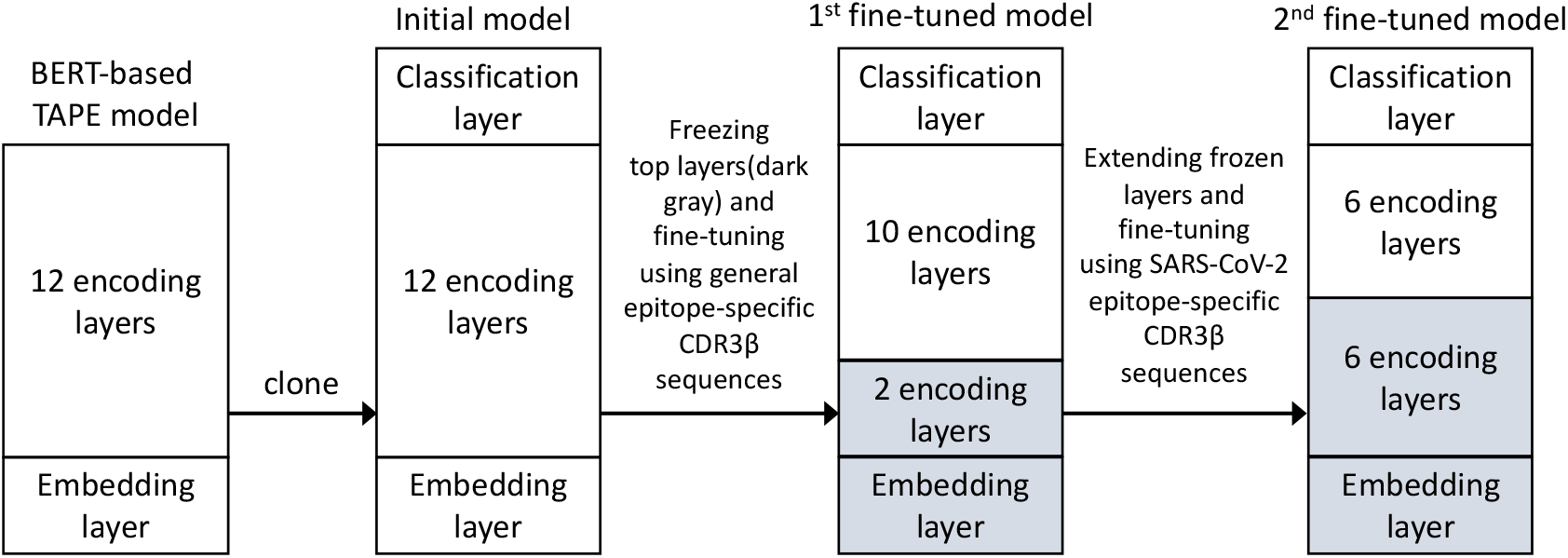
Training process for the proposed model. The initial model is cloned from pre-trained Tasks Assessing Protein Embeddings (TAPE) model, adding a classification layer at the end. The pre-trained model is fine-tuned in two rounds in a progressively specialized manner while extending the frozen layers between rounds.

Figure 2 shows the proposed model architecture. Input amino acid sequences concatenated by epitope and CDR3β sequences are first encoded into tokens using a tokenizer, where each token is an integer code for a single amino acid. Each token is then embedded into a 768-dimensional vector in the pre-trained TAPE model based on the BERT model which has 12 encoding layers with 12 self-attention heads in each layer. The TAPE model was pre-trained using 31 million unlabeled protein sequences, via next-token prediction and bidirectional masked-token prediction tasks, with further supervised training via protein-specific tasks, contact prediction and remote homology detection. The output of the pre-trained TAPE model is the hidden states of the first token. The final classifier, a 2-layer feed-forward network, is then used to predict either binder or not from the output of the TAPE model.

**Figure 2.**
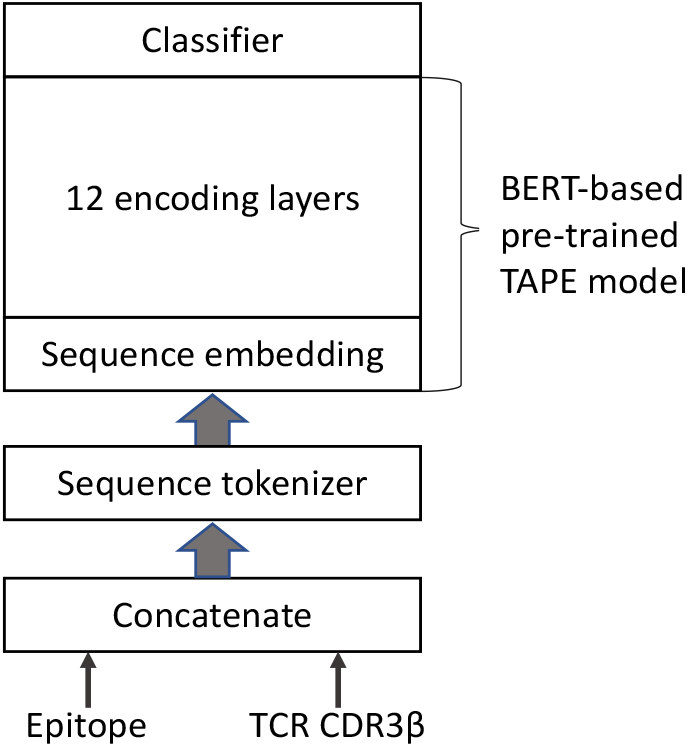
The proposed model architecture. Input amino acid sequences concatenated by epitope and CDR3β sequences are first encoded into tokens using a tokenizer. Each token is then embedded into a 768-dimensional vector in the pre-trained Tasks Assessing Protein Embeddings (TAPE) model which has 12 encoding layers with 12 self-attention heads in each layer. The final classifier, a 2-layer feed forward network, is then used to predict either binder or not from the output of the TAPE model.

### Datasets

#### Fine-tuning datasets

For the first fine-tuning round, the positive dataset containing epitope-specific TCR CDR3β sequences was compiled in May 2021 from three data sources: Dash *et al*.[Dash:2017go]^11^, providing epitope-specific paired TCRα and TCRβ chains for three human epitopes and seven mouse epitopes, and two manually curated databases providing pathology-associated TCR sequences, VDJdb [15] (https://vdjdb.cdr3.net) and McPAS-TCR [13] (http://friedmanlab.weizmann.ac.il/McPAS-TCR/). All VDJdb entries have confidence scores: 0, critical information missing; 1, medium confidence; 2, high confidence; 3, very high confidence. We selected all VDJdb entries with a confidence score of at least 1. For the second fine-tuning round, SARS-CoV-2 T-cell epitope-specific CDR3β sequence data were obtained from the Immune Epitope Database [35] (https://iedb.org) in June 2021. After selecting all epitopes with at least 20 CDR3β sequences and removing duplicates with the same combination of epitope and CDR3β sequences from each fine-tuning dataset, the datasets for the first and second fine-tuning rounds contained, respectively, 12,569 positive data points covering 78 epitopes, and 49,282 positive data points covering 145 epitopes.

To increase the specificity of our model, it was necessary to add more epitope-specific TCR CDR3β sequence data to each fine-tuning dataset as negative examples that are not expected to interact with TCRs and epitopes. Background CDR3β sequences were obtained from Howie *et al*. [36], who collected blood from two healthy donors. A negative example was generated by combining an epitope from the positive dataset and a randomly selected background TCR CDR3β sequence. Table S1 summarizes the final epitope-specific CDR3β sequence data for each fine-tuning dataset.

#### Evaluation datasets

We evaluated the final model using two independent datasets. The first dataset contained 305 COVID-19 **S**-protein_269-277_ T-cell epitope (**YLQPRTFLL**)-specific TCRβs from a recent study by Shomuradova *et al*. [37] (hereafter referred to as the Shomuradova dataset) and the same number of negative data points. The second dataset (hereafter referred to as the ImmuneCODE dataset) contained 390 **YLQPRTFLL**-specific TCRβs from the ImmuneRACE study launched on June 10, 2020, by Adaptive Biotechnologies and Microsoft (https://immunerace.adaptivebiotech.com), and 328 negative data points (Table S2).

### Finetuning and evaluating the model

We fine-tuned the pre-trained model in two rounds, changing the frozen layers between rounds in a progressively specialized manner. In the first fine-tuning round, the model was trained while freezing the embedding layer and top two encoding layers, so that the weights of the layers were not updated during the training process. In the second fine-tuning round, freezing was extended to the top six encoding layers. In each fine-tuning round, the training dataset was split into 80% training and 20% validation subsets, and training-validation was repeated for up to 200 epochs. The training and validation losses were measured for each epoch; the training process was stopped early at the epoch in which the validation loss had not been decreased for 15 consecutive epochs [38]. We used the Adam optimizer [39] with a learning rate of 0.0001 and batch size of 128, in all epochs. The PyTorch deep learning library(https://pytorch.org) was used to implement the model. We evaluated the final fine-tuned model using the Shomuradova and ImmuneCODE datasets and quantified its predictive performance using the area under a receiver operating characteristic (AUROC) score.

### Interpreting position-specific attention weights

To identify the critical amino acid positions in both the SARS-CoV-2 epitope (**YLQPRTFLL**) and CDR3β sequences, which potentially contribute greatly to TCR recognition of the epitope, we investigated the output attention weights of our model for the **YLQPRTFLL**-CDR3β sequence pairs predicted as a binder in the Shomuradova and ImmuneCODE datasets. We selected CDR3β sequences with the most common lengths of 13(n=159), 16(n=62), and 11(n=35) from the Shomuradova dataset, and 13(n=162), 14(n=60), and 16(n=58) from the ImmuneCODE dataset were selected. The output attention weights have the dimension (**L, N, H, S, S**), where **L** is the number of encoding layers, **N** is the number of **YLQPRTFLL**-CDR3β sequence pairs, **H** is the number of attention heads, and **S** is the fixed-length of the sequences. The attention weights were marginalized into a one-dimensional vector of length of **S**. A value of the vector at the position *m*, ***A***_***m***_ is given by the following equation:

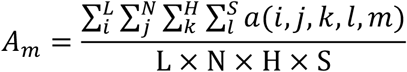

where, ***a(i, j, k, l, m)*** is an attention weight.

## Results and Discussion

### Finetuning results

The final validation accuracies were 0.793 and 0.934 in two fine-tuning rounds, respectively (Figure 3). In first fine-tuning round used a more general training dataset and more trainable encoding layers, the validation accuracy was lower and the difference between training and validation accuracies was higher. In contrast, in the second fine-tuning round used a more specific training dataset and fewer trainable encoding layers, the validation accuracy was markedly high and the difference between the training and validation accuracies was smaller. Fine-tuning the pre-trained model in this progressively specialized manner has the potential to generate a final model with high predictive performance for a specific task while avoiding model overfitting.

**Figure 3.**
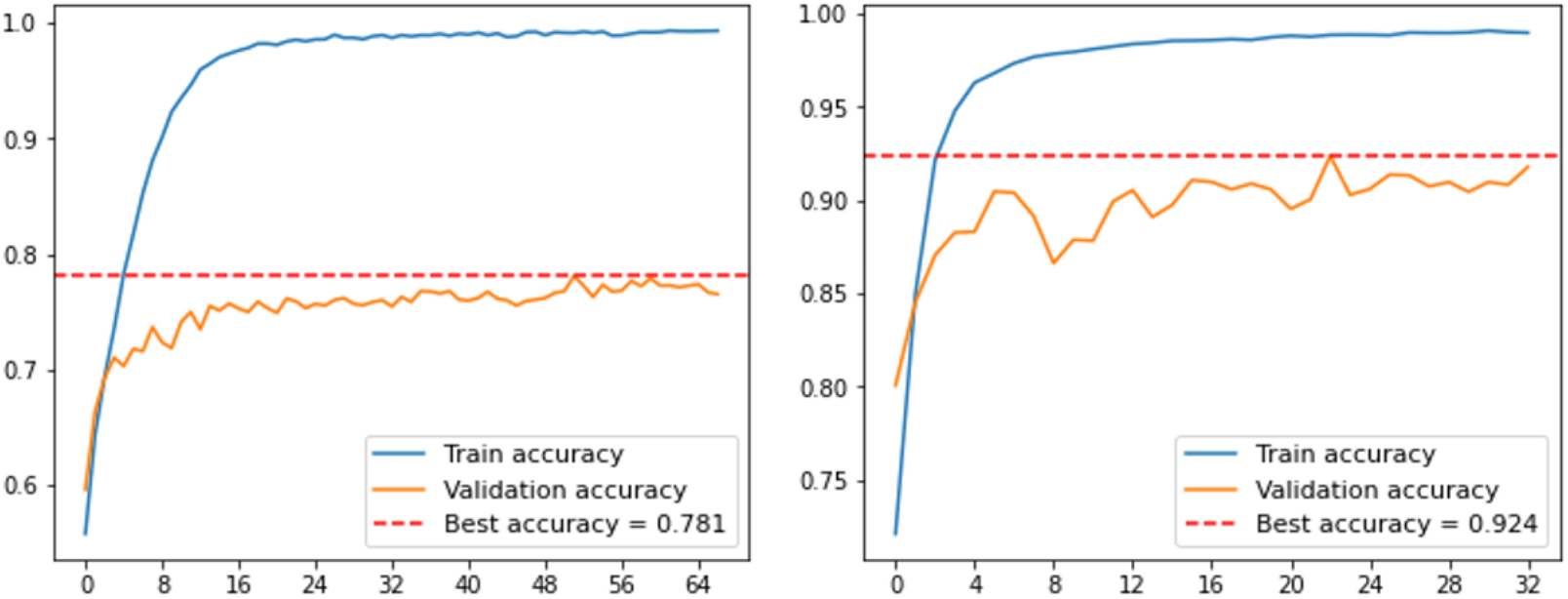
Fine-tuning of the pre-trained model in two rounds. The final validation accuracies were 0.781 and 0.924 in two fine-tuning rounds, respectively. Progressively, the validation accuracy was increased and the difference between the training and validation accuracies was reduced, in fine-tuning rounds.

### Evaluation results

The final fine-tuned model was evaluated using two external test datasets containing the SARS-CoV-2 epitope (**YLQPRTFLL**)-specific CDR3β sequences. Figure 4 shows the ROC curves for the two datasets. The AUROC scores were significantly high, at 0.981 and 0.983 for Shomuradova and ImmuneCODE datasets, respectively. Our model outperformed the recent Gaussian process-based method, TCRGP [19], which produced an AUROC score of 0.895 for the ImmuneCODE dataset.

**Figure 4.**
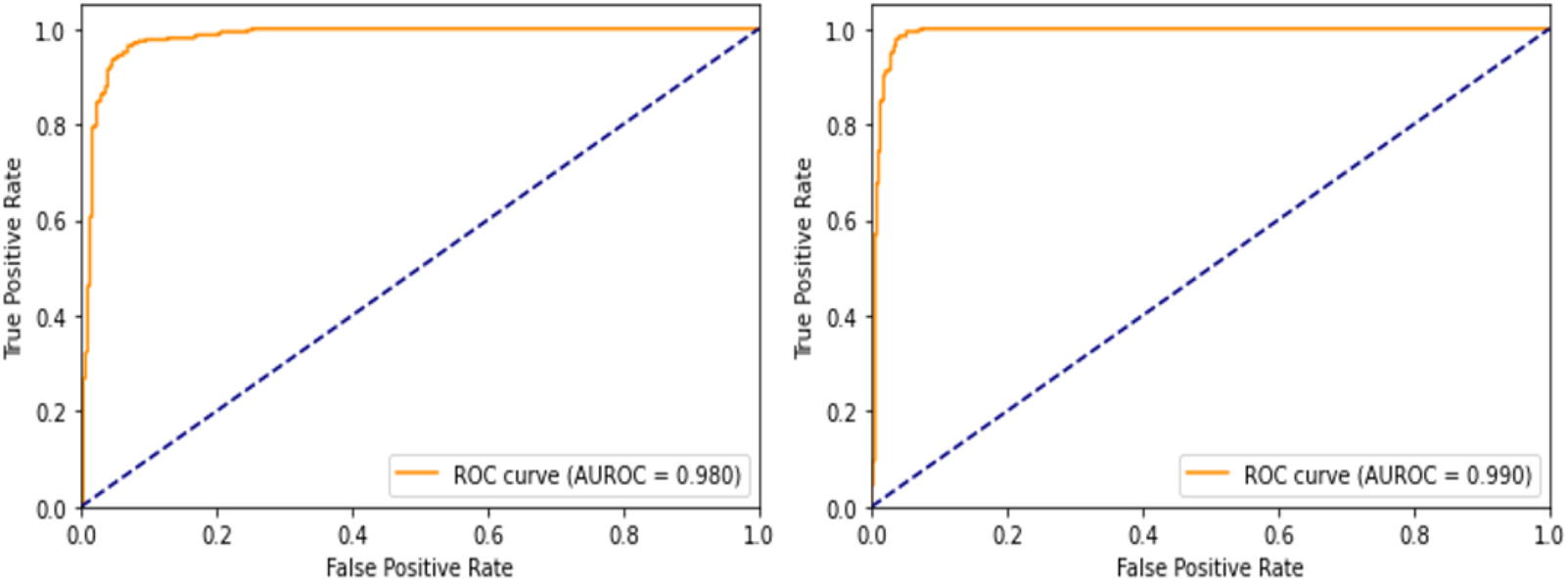
**Receiver operating characteristic (ROC) curves** for evaluating the final fine-tuned model using two external datasets containing the SARS-CoV-2 epitope (**YLQPRTFLL**)-specific CDR3β sequences. The area under the ROC (AUROC) scores were significantly high at 0.980 and 0.990 for the Shomuradova (A, left panel) and ImmuneCODE (B, right panel) datasets.

### Position-wise attention weight analysis

To identify critical amino acid positions in both **YLQPRTFLL** and CDR3β sequences, we investigated the output attention weights of our model for the **YLQPRTFLL**-CDR3β sequence pairs predicted as a binder from the Shomuradova and ImmuneCODE datasets (Figure 5).

**Figure 5.**
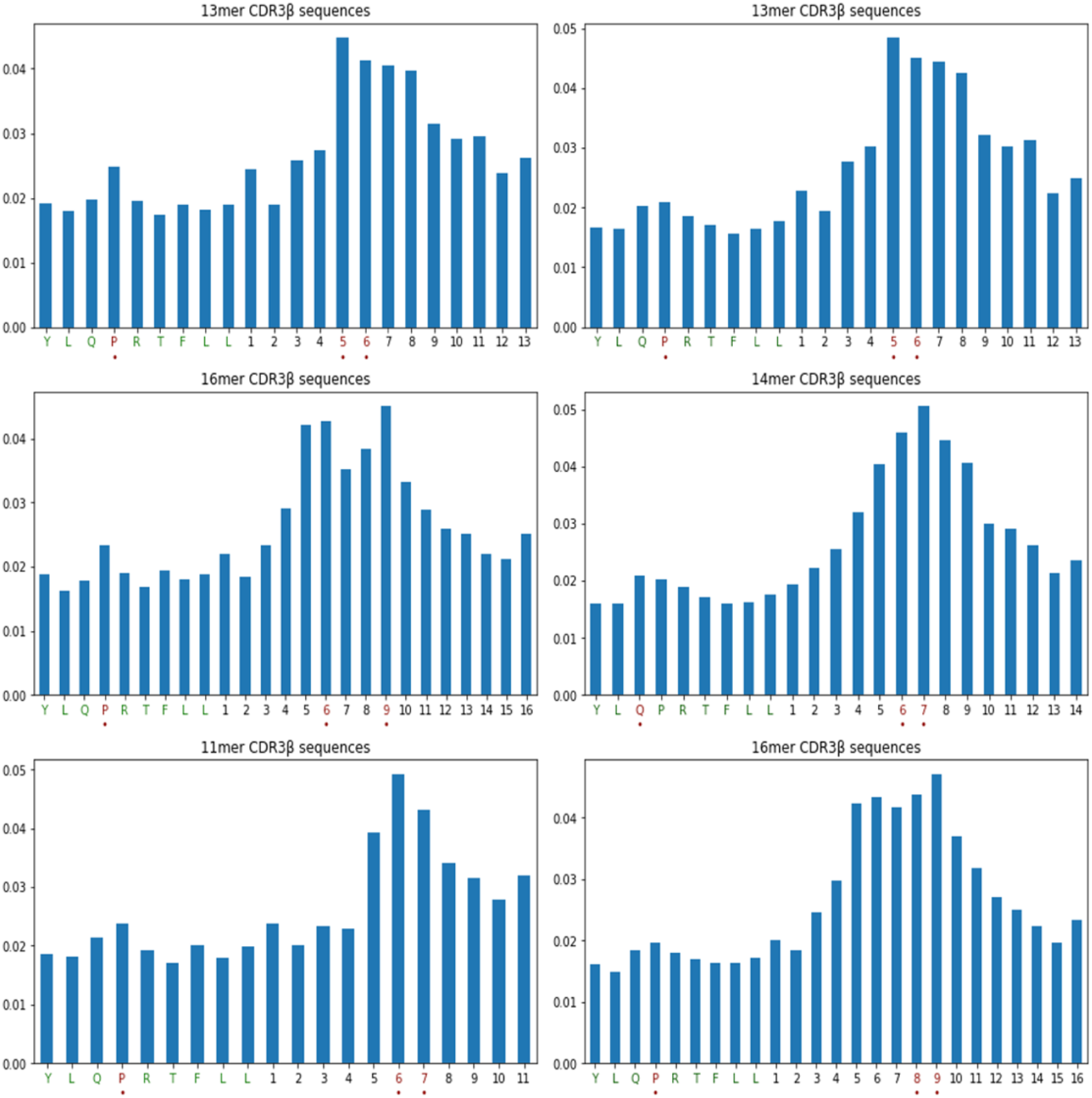
**Marginalized position-wise attention weights** for the **YLQPRTFLL**-CDR3β sequence pairs predicted as a binder from the Shomuradova (A, left panels) and ImmuneCODE (B, right panels) datasets. The CDR3β sequence lengths differ from top to bottom. The amino acid positions corresponding to the top 10% weights of each the epitope and CDR3 sequences are highlighted in red dots below x-axis ticks.

For the Shomuradova dataset, the proline at the position 4 (P4) in the epitope has a relatively high attention weight, indicating that P4 may have a critical contribution to TCR recognition of the epitope (Figure 5A). A recent experimental study [40] of SARS-CoV-2 variants found that CD8^+^ T-cells from a cohort of convalescent patients, comprising more than 120 different TCRs, failed to respond to the **P272L** variant corresponding to P4. Furthermore, sizable populations of CD8^+^ T cells from individuals immunized with the currently approved COVID-19 vaccines failed to bind to the **P272L** reagent. In the CDR3β sequences, the attention weights at the central positions 5-8 were higher than those at both ends, indicating that the TCR amino acids at the positions may interact relatively well with the proline at P4 of the epitope, thereby contributing substantially to TCR recognition of the epitope.

Very similar attention weight patterns were observed for the ImmuneCODE dataset (Figure 5B), for both the epitope and CDR3β sequences: there were relatively high attention weights at P4 in the epitope and the central positions 6-9 in the CDR3β sequences. Interestingly, our attention-based results differed from those of the MSA-based approaches, which consider conserved positions to be highly relevant to epitope-specific TCR recognition. In contrast, our findings suggest that the variable amino acid positions in the CDR3β sequences contribute substantially to TCR recognition of the epitope (Figure S1 provides the sequence logos [41] of MSAs of the CDR3β sequences).

## Conclusion

We developed a BERT-based model employing self-supervised transfer learning to predict SARS-CoV-2 epitope-specific TCR recognition. The predictive model was generated by fine-tuning the pre-trained TAPE model using epitope-specific TCR CDR3β sequence datasets in a progressively specialized manner. The fine-tuned model demonstrated a markedly high predictive performance for two evaluation datasets containing the SARS-CoV-2 **S**-protein_269-277_ epitope (**YLQPRTFLL**)-specific CDR3β sequences, and outperformed the recent Gaussian process-based model, TCRGP, for the ImmuneCODE dataset. In particular, the output attention weights of our model suggest that the proline at **P4** in the epitope may contribute critically to TCR recognition of the epitope. A recent experimental study of SARS-CoV-2 variants demonstrated that CD8^+^ T-cells failed to respond to the **P272L** variant corresponding to P4. Further, CDR3β-sequence amino acids at the central positions, rather than at both ends, may contribute to the TCR recognition of the epitope. Our attention-based approach, which can capture all motifs in both the epitope and CDR3β sequences in epitope-specific TCR recognition, may be more useful for predicting immunogenic changes in T-cell epitopes derived from SARS-CoV-2 mutations than MSA-based approaches which depend entirely on TCR sequences.

In further studies, sequence data related to interactions between TCRα chains and MHC molecules will be integrated into our framework to predict global interaction patterns in TCR recognition of peptide-MHC complexes. We anticipate that our findings will provide new frameworks for constructing a reliable data-driven model for predicting the immunogenic T cell epitopes using limited training data and help accelerate the development of an effective vaccine for the response to SARS-CoV-2 variants, by identifying critical amino acid positions that are important in epitope-specific TCR recognition.

## Supporting information

All figures

## Acknowledgements

The authors would like to thank Dr. G. Kim and W. Jeon for helpful discussions and comments.

## Funding

This work was supported by the research program of Korea Institute of Science and Technology Information (KISTI).

## Availability of data and materials

Python source codes and all the datasets supporting this work can be downloaded from https://github.com/luseedbio/TCRBert.

## Authors’ contributions

YH designed the method, developed the software, conducted the experiments, and wrote the manuscript. AL supported the experiments and writing. All authors read and approved the final manuscript.

## Competing interests

The authors declare that they have no competing interests.

## Notes

### Competing Interest Statement

The authors have declared no competing interest.

## References

1. Elbe, S. & Buckland-Merrett, G. Data, disease and diplomacy: GISAID’s innovative contribution to global health. Global Challenges 1, 33–46 (2017).

2. Tao, K. et al. The biological and clinical significance of emerging SARS-CoV-2 variants. Nat Rev Genet 1–17 (2021) doi:10.1038/s41576-021-00408-x.

3. Oh, H.-L. J., Gan, S. K.-E., Bertoletti, A. & Tan, Y.-J. Understanding the T cell immune response in SARS coronavirus infection. Emerging Microbes & Infections 1, 1–6 (2019).

4. Channappanavar, R., Zhao, J., research, S. P. I. & 2014. T cell-mediated immune response to respiratory coronaviruses. Immunologic research 118–128 (2014).

5. Channappanavar, R. et al. Virus-Specific Memory CD8 T Cells Provide Substantial Protection from Lethal Severe Acute Respiratory Syndrome Coronavirus Infection. Journal of Virology 88, 11034–11044 (2014).

6. Yang, L. et al. Persistent memory CD4+ and CD8+ T-cell responses in recovered severe acute respiratory syndrome (SARS) patients to SARS coronavirus M antigen. The Journal of general virology 88, 2740–2748 (2007).

7. Rudolph, M. G., Stanfield, R. L. & Wilson, I. A. HOW TCRS BIND MHCS, PEPTIDES, AND CORECEPTORS. Annu Rev Immunol 24, 419–466 (2006).

8. Bassing, C. H., Swat, W. & Alt, F. W. The Mechanism and Regulation of Chromosomal V(D)J Recombination. Cell 109, S45–S55 (2002).

9. Robins, H. S. et al. Comprehensive assessment of T-cell receptor β-chain diversity in αβ T cells. Blood 114, 4099–4107 (2009).

10. Glanville, J. et al. Identifying specificity groups in the T cell receptor repertoire. Nature 547, 94–98 (2017).

11. Dash, P. et al. Quantifiable predictive features define epitope-specific T cell receptor repertoires. Nature 547, 89–93 (2017).

12. Borrman, T. et al. ATLAS: A database linking binding affinities with structures for wild-type and mutant TCR-pMHC complexes. Proteins: Structure, Function, and Bioinformatics 85, 908–916 (2017).

13. Tickotsky, N., Sagiv, T., Prilusky, J., Shifrut, E. & Friedman, N. McPAS-TCR: a manually curated catalogue of pathology-associated T cell receptor sequences. Bioinformatics 33, 2924–2929 (2017).

14. Mahajan, S. et al. Epitope Specific Antibodies and T Cell Receptors in the Immune Epitope Database. Frontiers in Immunology 9, 3628–10 (2018).

15. Bagaev, D. V. et al. VDJdb in 2019: database extension, new analysis infrastructure and a T-cell receptor motif compendium. Oxford University Press 1–6 (2019) doi:10.1093/nar/gkz874.

16. Klinger, M. et al. Multiplex Identification of Antigen-Specific T Cell Receptors Using a Combination of Immune Assays and Immune Receptor Sequencing. PLoS ONE 10, e0141561 (2015).

17. Bentzen, A. K. et al. Large-scale detection of antigen-specific T cells using peptide-MHC-I multimers labeled with DNA barcodes. Nature Biotechnology 34, 1037–1045 (2016).

18. Zvyagin, I. V., Tsvetkov, V. O., Chudakov, D. M. & Shugay, M. An overview of immunoinformatics approaches and databases linking T cell receptor repertoires to their antigen specificity. Immunogenetics 1–8 (2019) doi:10.1007/s00251-019-01139-4.

19. Jokinen, E., Huuhtanen, J., Mustjoki, S., Heinonen, M. & Lähdesmäki, H. Predicting recognition between T cell receptors and epitopes with TCRGP. PLoS Computational Biology 17, e1008814 (2021).

20. Gielis, S. et al. Detection of Enriched T Cell Epitope Specificity in Full T Cell Receptor Sequence Repertoires. Frontiers in Immunology 10, 2820 (2019).

21. Jurtz, V. I. et al. NetTCR: sequence-based prediction of TCR binding to peptide-MHC complexes using convolutional neural networks. bioRxiv 433706 (2018) doi:10.1101/433706.

22. Isacchini, G., Walczak, A. M., Mora, T. & Nourmohammad, A. Deep generative selection models of T and B cell receptor repertoires with soNNia. Proceedings of the National Academy of Sciences 118, (2021).

23. Sidhom, J.-W., Larman, H. B., Pardoll, D. M. & Baras, A. S. DeepTCR is a deep learning framework for revealing sequence concepts within T-cell repertoires. Nature Communications 1–12 (2021) doi:10.1038/s41467-021-21879-w.

24. Springer, I., Besser, H., Tickotsky-Moskovitz, N., Dvorkin, S. & Louzoun, Y. Prediction of Specific TCR-Peptide Binding From Large Dictionaries of TCR-Peptide Pairs. Frontiers in Immunology 11, 1803 (2020).

25. Peters, M. E. et al. Deep contextualized word representations. http://arXiv.org cs.CL, (2018).

26. Devlin, J., Chang, M.-W., Lee, K. & Toutanova, K. BERT: Pre-training of Deep Bidirectional Transformers for Language Understanding. http://arXiv.org cs.CL, (2018).

27. Radford, A., Wu, J. & Child, R. Language models are unsupervised multitask learners. OpenAI blog (2019).

28. Rao, R. et al. Evaluating Protein Transfer Learning with TAPE. Advances in neural information processing systems 32, 9689–9701 (2019).

29. Heinzinger, M. et al. Modeling aspects of the language of life through transfer-learning protein sequences. BMC Bioinformatics 20, 1–17 (2019).

30. Nambiar, A. et al. Transforming the language of life: transformer neural networks for protein prediction tasks. Proceedings of the 11th ACM International Conference on Bioinformatics, Computational Biology and Health Informatics, 1–8 (2020).

31. El-Gebali, S. et al. The Pfam protein families database in 2019. Nucleic Acids Res 47, gky995.(2018).

32. Cheng, J., Bendjama, K., Rittner, K. & Malone, B. BERTMHC: Improves MHC-peptide class II interaction prediction with transformer and multiple instance learning. bioRxiv 2020.11.24.396101 (2020) doi:10.1101/2020.11.24.396101.

33. Vig, J. et al. BERTology Meets Biology: Interpreting Attention in Protein Language Models. Arxiv cs.CL, (2020).

34. Jin, J. et al. Deep learning pan-specific model for interpretable MHC-I peptide binding prediction with improved attention mechanism. Proteins Struct Funct Bioinform 89, 866–883 (2021).

35. Vita, R. et al. The immune epitope database (IEDB) 3.0. Nucleic Acids Research 43, D405–12 (2015).

36. Howie, B. et al. High-throughput pairing of T cell receptor α and β sequences. Science Translational Medicine 7, 301ra131–301ra131 (2015).

37. Shomuradova, A. S. et al. SARS-CoV-2 Epitopes Are Recognized by a Public and Diverse Repertoire of Human T Cell Receptors. Immunity 53, 1245–1257.e5 (2020).

38. Prechelt, Lutz. Early stopping-but when? Neural Networks: Tricks of the trade. Springer, Berlin, Heidelberg, 55–69 (1998).

39. Kingma, D. P. & Ba, J. Adam: A Method for Stochastic Optimization. http://arXiv.org cs.LG, (2014).

40. Dolton, G. et al. Emergence of immune escape at dominant SARS-CoV-2 killer T-cell epitope. medRxiv, (2021).

41. Thomsen, M. C. F. & Nielsen, M. Seq2Logo: a method for construction and visualization of amino acid binding motifs and sequence profiles including sequence weighting, pseudo counts and two-sided representation of amino acid enrichment and depletion. Nucleic Acids Research 40, W281–W287 (2012).

