## Supplementary material for "Predicting SARS-CoV-2 epitope-specific TCR recognition using pre-trained protein embeddings": All figures

### Slide 1
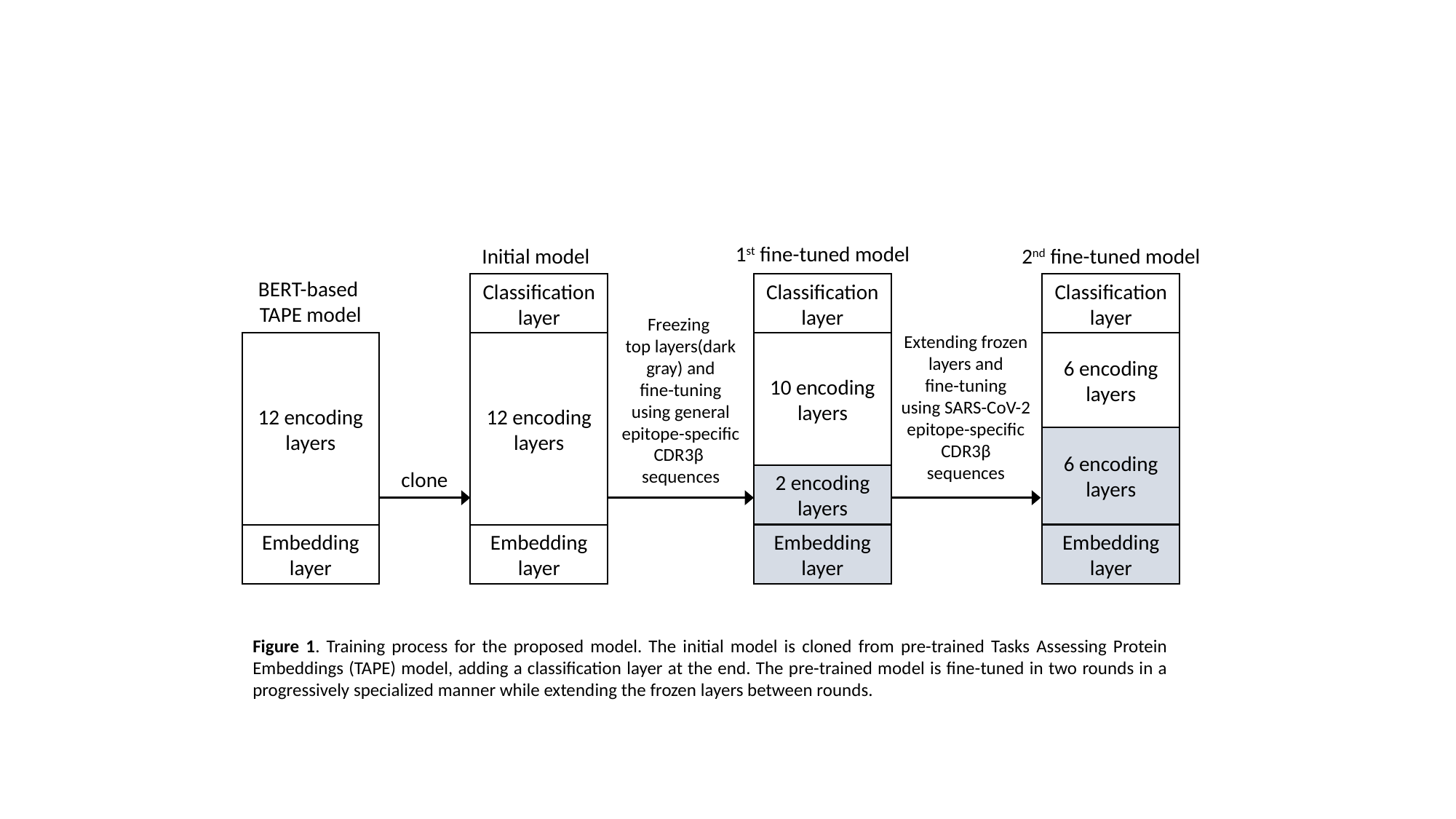

1^st^ fine-tuned model
Classification layer
10 encoding layers
2 encoding layers
Embedding layer
2^nd^ fine-tuned model
Classification layer
6 encoding layers
6 encoding layers
Embedding layer
Initial model
BERT-based
TAPE model
Classification layer
12 encoding layers
Embedding layer
Freezing
top layers(dark gray) and
fine-tuning
using general
epitope-specific CDR3β sequences
Extending frozen layers and
fine-tuning
using SARS-CoV-2
epitope-specific CDR3β sequences
12 encoding layers
Embedding layer
clone
Figure 1. Training process for the proposed model. The initial model is cloned from pre-trained Tasks Assessing Protein Embeddings (TAPE) model, adding a classification layer at the end. The pre-trained model is fine-tuned in two rounds in a progressively specialized manner while extending the frozen layers between rounds.

### Slide 2
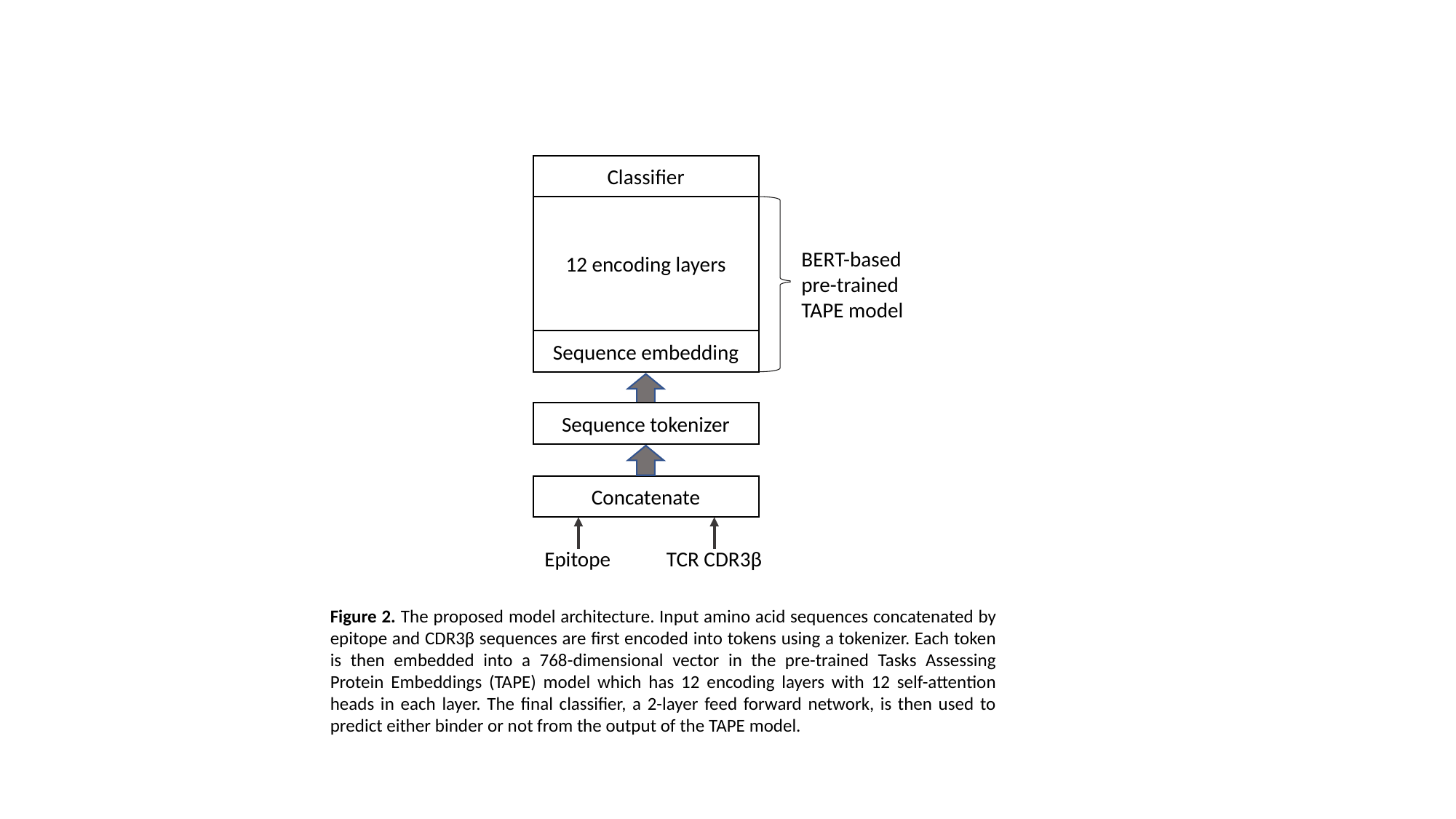

Classifier
12 encoding layers
Sequence embedding
BERT-based pre-trained TAPE model
Sequence tokenizer
Concatenate
Epitope
TCR CDR3β
Figure 2. The proposed model architecture. Input amino acid sequences concatenated by epitope and CDR3β sequences are first encoded into tokens using a tokenizer. Each token is then embedded into a 768-dimensional vector in the pre-trained Tasks Assessing Protein Embeddings (TAPE) model which has 12 encoding layers with 12 self-attention heads in each layer. The final classifier, a 2-layer feed forward network, is then used to predict either binder or not from the output of the TAPE model.

### Slide 3
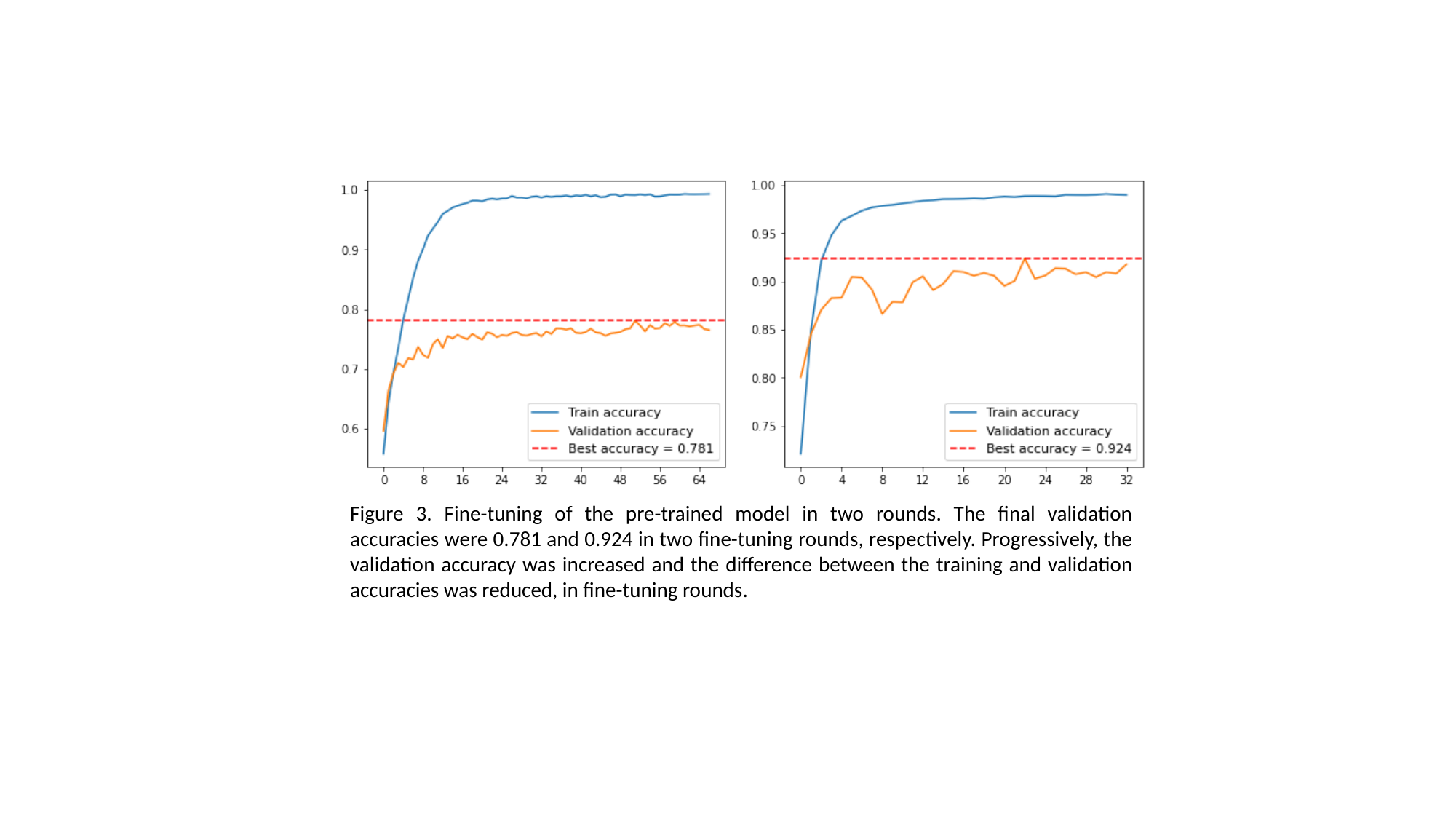

Figure 3. Fine-tuning of the pre-trained model in two rounds. The final validation accuracies were 0.781 and 0.924 in two fine-tuning rounds, respectively. Progressively, the validation accuracy was increased and the difference between the training and validation accuracies was reduced, in fine-tuning rounds.

### Slide 4
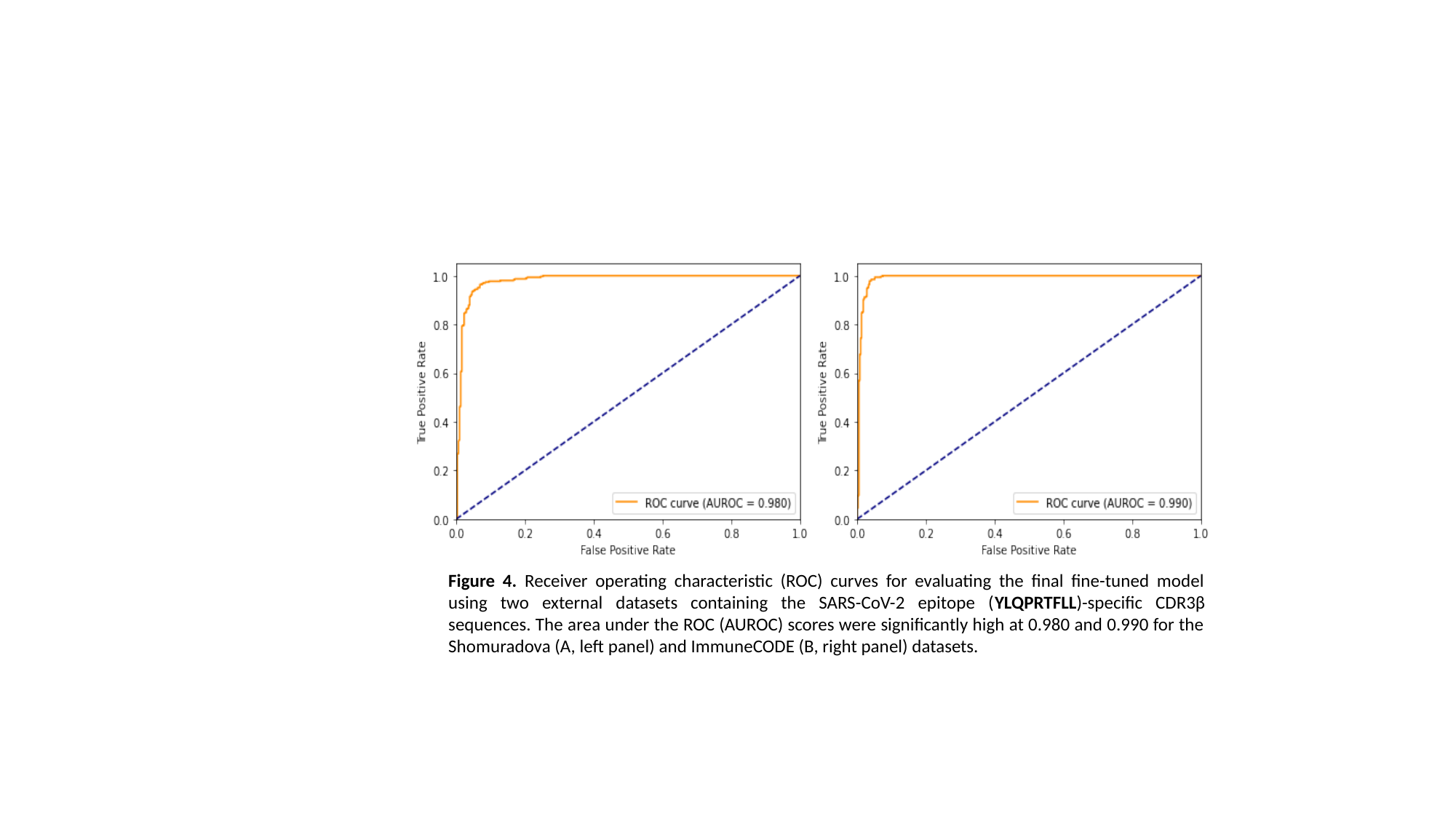

Figure 4. Receiver operating characteristic (ROC) curves for evaluating the final fine-tuned model using two external datasets containing the SARS-CoV-2 epitope (YLQPRTFLL)-specific CDR3β sequences. The area under the ROC (AUROC) scores were significantly high at 0.980 and 0.990 for the Shomuradova (A, left panel) and ImmuneCODE (B, right panel) datasets.

### Slide 5
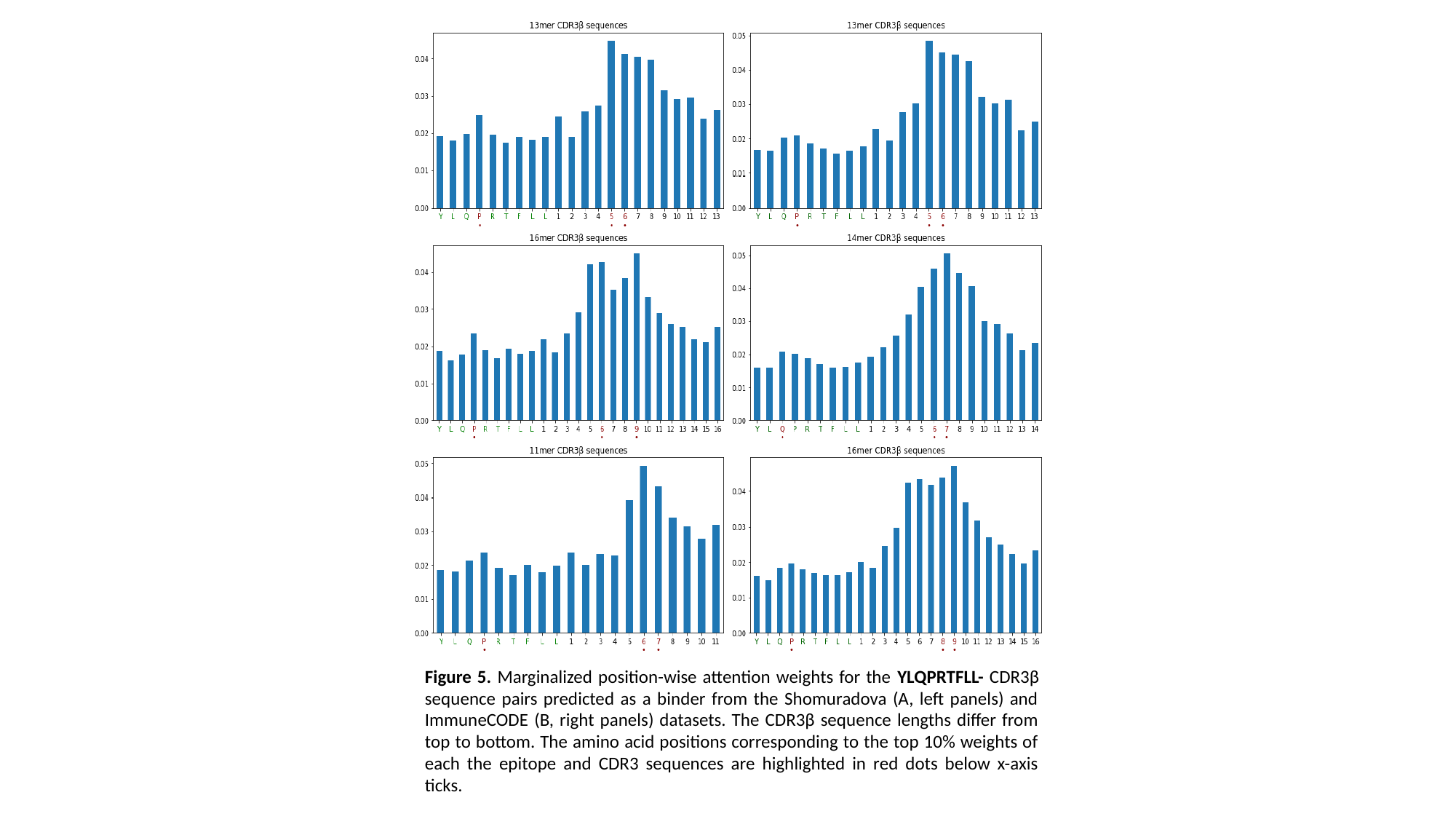

Figure 5. Marginalized position-wise attention weights for the YLQPRTFLL- CDR3β sequence pairs predicted as a binder from the Shomuradova (A, left panels) and ImmuneCODE (B, right panels) datasets. The CDR3β sequence lengths differ from top to bottom. The amino acid positions corresponding to the top 10% weights of each the epitope and CDR3 sequences are highlighted in red dots below x-axis ticks.

### Slide 6
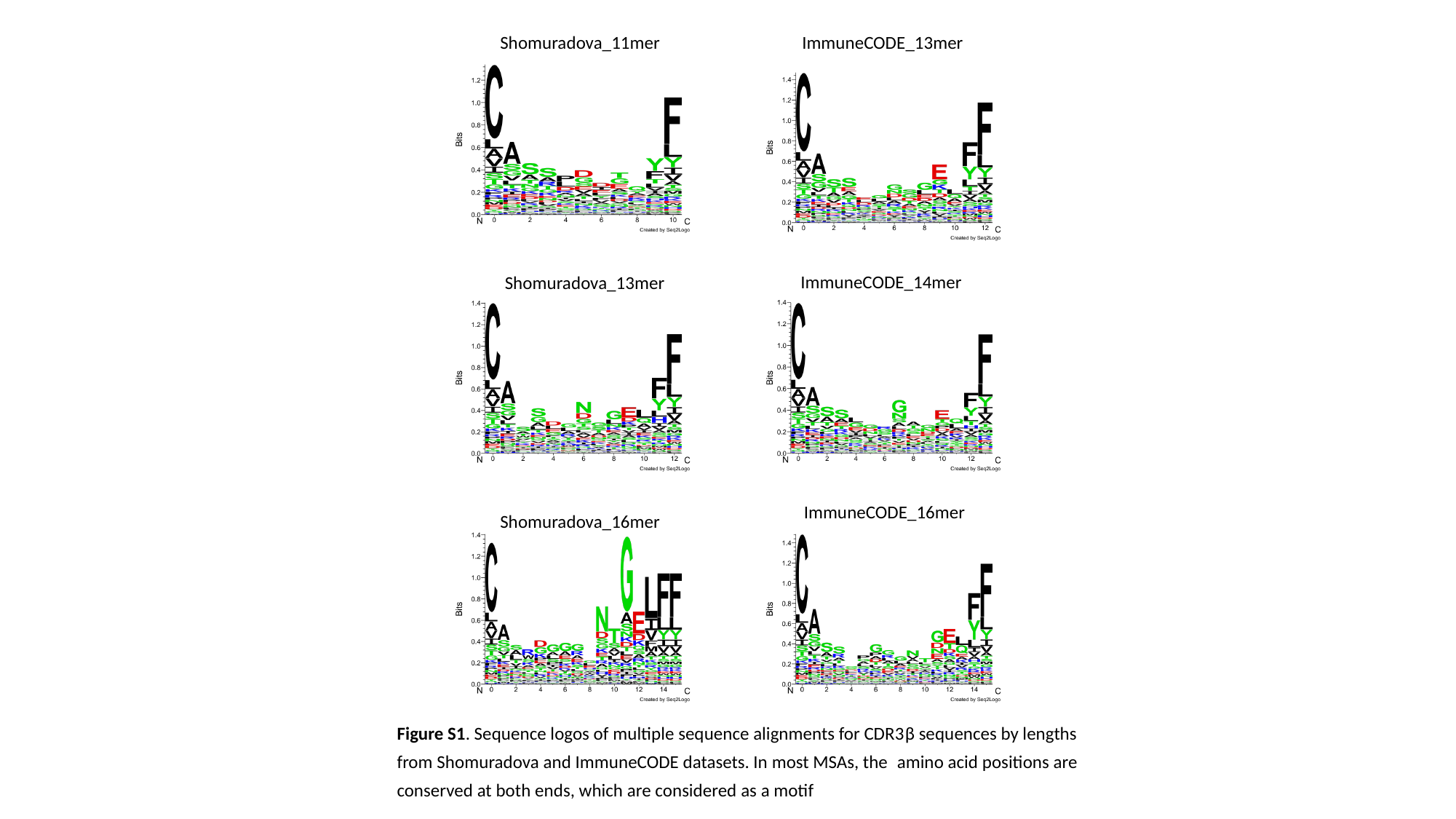

Shomuradova_11mer
ImmuneCODE_13mer
ImmuneCODE_14mer
Shomuradova_13mer
ImmuneCODE_16mer
Shomuradova_16mer
Figure S1. Sequence logos of multiple sequence alignments for CDR3β sequences by lengths from Shomuradova and ImmuneCODE datasets. In most MSAs, the amino acid positions are conserved at both ends, which are considered as a motif
